# Multivariate Resting-State Functional Connectivity Features Linked to Transdiagnostic Psychopathology in Early Psychosis

**DOI:** 10.1101/2025.06.04.654984

**Authors:** Haley R. Wang, Zhen-Qi Liu, Jason S. Nomi, Charles H. Schleifer, Carrie E. Bearden, Bratislav Misic, Lucina Q. Uddin, Katherine H. Karlsgodt

## Abstract

**Background:** Early psychosis (EP) is characterized by neurobiological changes, including alterations in resting-state functional connectivity (RSFC). We now understand that symptoms and neural changes may overlap across EP diagnostic categories. However, the relationship between RSFC patterns and transdiagnostic symptom dimensions remains poorly understood.

**Methods:** We employed Partial Least Squares correlation to examine multivariate relationships between whole-brain RSFC and clinical symptoms in 124 EP patients (aged 16-35 years) diagnosed with schizophrenia, schizoaffective disorder, or a psychotic mood disorder. RSFC was computed among 216 cortical and subcortical regions. Clinical assessment included 41 symptom measures spanning positive, negative, general psychopathology, and manic dimensions.

**Results:** Analysis revealed one significant latent component (p<0.001) capturing 41.6% of the RSFC-symptom covariance. This component was characterized by increased between-network connectivity, particularly involving sensory-motor, default mode, and subcortical regions including the amygdala and thalamus. The associated symptom profile included cognitive rigidity and arousal dysregulation (stereotyped thinking, anxiety, and somatic concerns), rather than traditional positive or negative symptoms. This brain-behavior relationship was consistent across diagnoses and independent of medication and substance use. The clinical relevance was validated through significant correlations with standardized measures of hostility (r=0.23), negative affect (r=0.25), and perceived stress (r=0.22).

**Conclusions:** Our findings reveal a distinct transdiagnostic phenotype in EP characterized by cognitive inflexibility and arousal dysregulation that is associated with altered integration between sensory, default mode, and subcortical networks. This work suggests that specific patterns of network-level functional connectivity may relate to symptom dimensions that cut across conventional diagnostic boundaries, potentially informing more targeted therapeutic approaches.

## Introduction

Early psychosis (EP) refers to the first three years following the onset of psychotic symptoms, and encompasses diagnoses such as schizophrenia, schizoaffective disorder, and bipolar disorder with psychotic features (1, 2). This critical period is characterized by significant overlapping symptoms across diagnostic categories, diagnostic instability, and substantial neurobiological changes that represent both transdiagnostic vulnerability markers and dynamic correlates of symptom severity (3–5). Understanding the shared neurobiological mechanisms underlying psychotic disorders during this period is essential for developing targeted interventions for youth with EP, yet remains a significant challenge in current research.

Resting-state functional connectivity (RSFC), which putatively represents the intrinsic functional organization of brain regions that support affective and cognitive processes, has been found to be disrupted in psychotic spectrum disorders (6–8). Extensive resting-state fMRI meta- and mega-analyses have revealed widespread RSFC reductions in individuals with psychosis compared to controls, with the most substantial effects observed in the default mode network (medial prefrontal cortex, posterior cingulate/precuneus), the salience network (anterior insula, dorsal anterior cingulate), and the central executive network (dorsolateral prefrontal cortex, posterior parietal cortex) (4, 9, 10). While these RSFC alterations have been associated with symptom severity and cognitive deficits in schizophrenia and related disorders, findings remain inconsistent and investigations are often confined to specific diagnostic categories (11–13).

The patterns of RSFC disruption observed in psychosis show both shared and distinct features across diagnoses. In schizophrenia, large-scale meta-analyses reveal a characteristic pattern: hypoconnectivity within higher-order associative networks (default mode, salience, frontoparietal control) accompanied by hyperconnectivity in subcortical and sensorimotor circuits (e.g., thalamus–sensorimotor and insula–striatal pathways), consistently implicating nodes such as the insula, lateral postcentral cortex, striatum, and thalamus (4, 6, 9, 12). In contrast, bipolar disorder with psychotic features and schizoaffective disorder typically demonstrate decreased RSFC between limbic hubs (amygdala, ventral striatum) and medial prefrontal cortex, alongside dysconnectivity in dorsal and ventral attention networks (14–19). Both schizophrenia and psychotic mood disorders share hypoconnectivity in frontoparietal control and default mode networks—reflecting common impairments in executive control and self-referential processing (8, 20). However, schizophrenia more consistently exhibits hyperconnectivity in thalamus–sensorimotor and insula–striatal circuits, whereas psychotic mood disorders show greater limbic–prefrontal coupling deficits (e.g., amygdala–ventromedial PFC) and attentional network alterations (16).

Despite these valuable insights into disorder-specific neural alterations, few studies have explicitly examined shared neurobiological underpinnings across diagnoses. Studies that have explored this (12, 21) suggest that covariance between brain and behavior in psychotic illness may be shared across affective and non-affective psychosis, though comprehensive symptom patterns remain unexplored. For example, EP patients show consistent reductions in within-network connectivity in major cortical systems—most notably the default mode and frontoparietal control networks—relative to healthy controls, regardless of specific diagnosis (22, 23). Given EP’s diagnostic fluidity and variable prognosis, identifying shared neurobiological correlates of symptom dimensions—rather than relying on categorical diagnoses—may offer greater clinical utility for understanding illness progression (24). Given adolescence-specific neurodevelopmental alterations—including intensified synaptic pruning, ongoing myelination, and functional connectivity reorganization—diagnostic fluidity during adolescence may reflect variable trajectories of vulnerability (25), suggesting that symptom dimensions could provide greater clinical insight than categorical diagnoses during this developmental window (5, 26).

Traditional brain-behavior correlation studies in psychosis research have predominantly relied on univariate methods, examining one-to-one relationships between neural connectivity and specific symptom measures. For instance, negative symptoms have been correlated with cerebellar-cerebral functional connectivity (27), and specific network relationships have been correlated with various symptom scores in bipolar and schizophrenia patients (16). Though informative, these theory-driven approaches have varied in factors such as symptom selection and RSFC regions of interest, making it challenging to synthesize findings into a cohesive understanding and limiting exploration of more complex, multivariate RSFC-psychopathology correlations. In contrast, data driven multivariate approaches like Partial Least Squares (PLS) correlation offer distinct advantages for investigating brain-behavior relationships. PLS can identify complex optimally covarying patterns between high-dimensional brain features such as RSFC and clinical features (1, 12, 28) by revealing latent components (LCs) representing key brain-behavior variable patterns. PLS is also able to manage collinearity and enables flexible post-hoc analyses of LCs associations (29, 30). Neurobiologically, PLS facilitates a nuanced, feature-level understanding of how various brain metrics, including whole-brain RSFC patterns, may interrelate with symptoms. Clinically, PLS allows for the organic emergence of symptom clusters that may not align with traditional groupings but potentially share underlying neurobiological mechanisms (1, 30). By accounting for EP’s clinical heterogeneity, PLS has the potential to uncover novel insights into the complex relationship between brain function and psychotic symptomatology, while shifting research from single diagnostic models to a transdiagnostic perspective.

In this study, we utilize PLS correlation to investigate the relationship between RSFC alterations and psychopathology dimensions in a transdiagnostic EP sample. Additionally, we examine the stability of the identified RSFC signature by accounting for potential confounders, including antipsychotic medication load, substance use, and illness duration. To validate the clinical utility of our findings, we associate the PLS-detected symptom clusters with external behavioral traits and psychosocial stressor measures. This comprehensive and transdiagnostic approach aims to provide a deeper understanding of the shared neurobiological mechanisms underlying psychotic disorders, while also bridging the gap between brain function, psychopathology, and behavioral traits.

## Methods and Materials

### Sample characteristics

The Human Connectome Project Early Psychosis Release 1.1 (HCP-EP) study cohort comprises 124 patients aged 16-35 years (mean 22.7 ± 3.7 years; 38.7% female). Participants had been diagnosed with schizophrenia, schizoaffective disorders, or psychotic mood disorders within the past three years. A healthy control group (n = 58) from the HCP-EP study was included for a supplementary comparison analysis. Table 1 provides a comprehensive overview of the cohort’s demographic and clinical characteristics. For detailed recruitment information, refer to Supplemental Methods 1-4. Figure S1 illustrates the differences in clinical scores across diagnostic groups.

**Table 1.** Demographic, clinical, and medication characteristics of the HCP-EP study participants.

| Characteristics | EP (n=124) | Controls (n=58) | P Value |
| --- | --- | --- | --- |
| Age, mean (SD), y | 22.7 (3.7) | 24.8 (4.1) | <0.001 |
| Male, No. (%) | 76 (61.3) | 38 (65.5) | 0.7 |
| Race, No. (%) |  |  |  |
| Asian | 8 (6.5) | 9 (14.8) | <0.001 |
| African American | 49 (39.5) | 5 (8.6) |  |
| American Indian/Alaska Native | 1 (0.8) | 0 (0) |  |
| White | 62 (50) | 42 (72.4) |  |
| Unknown or more than one race | 4 (3.2) | 3 (5.2) |  |
| Hispanic ethnicity, No. (%) | 8 (6.5) | 9 (15.5) | 0.09 |
| Parental education, mean (SD), y | 11.4 (2.5) | 11.7 (2.2) | 0.48 |
| WASI Full Scale IQ, mean (SD) | 101.4 (17.0) | 116 (10.7) | <0.001 |
| Duration of Illness, mean (SD), y | 2.19 (1.4) | NA | NA |
| Cannabis use in past 30 days, No. (%) | 16 (12.9) | NA | NA |
| Medication Use |  |  |  |
| Current dosage of chlorpromazine equivalent, mean (SD), mg | 159 (224.5) | NA | NA |
| SCID diagnosis, No. (%) |  |  |  |
| Schizophrenia | 81 (65.3) | NA | NA |
| Schizoaffective Disorder | 14 (11.3) |  |  |
| Bipolar I Disorder or Major Depressive Disorder with psychotic features | 26 (21) |  |  |
| YMRS score, mean (SD) | 4.9 (5.3) | NA | NA |
| PANSS scores, mean (SD) |  |  |  |
| Positive | 11.2 (4) | NA | NA |
| Negative | 14 (5.4) |  |  |
| General | 24.7 (5.2) |  |  |

### RSFC features

Rs-fMRI data was preprocessed using standard pipelines (31, 32) (Supplementary Methods 5-6). Three participants were excluded due to missing structural scans. Preprocessing included visual inspection for excessive motion and ICA-AROMA for minimizing motion-related artifacts (33). Because the global signal has been shown to correlate with key clinical and behavioral measures in psychosis— such that its removal could eliminate genuine disease□related variance—we did not apply global signal regression in our main analysis (34–36). However, acknowledging the disputed status of global signal regression (37, 38), we performed a secondary analysis with global signal regressed out from RSFC before PLS analysis for robustness control (Supplemental Methods 7).

RSFC was computed using Pearson’s correlation among the average time series of both cortical and subcortical regions. Cortical regions were identified using the 200 regions of interest (ROI) Schaefer 2018 atlas parcellation (39), which maps onto 17 distinct networks (Figure S2A). Subcortical regions included 16 parcels from a brain atlas developed by Tian et al. (2020) (Figure S2B) (40). Together covering the entire brain, the combined cortical and subcortical parcellation resulted in a 216 x 216 RSFC matrix for each participant. Site harmonization was performed on parcel-level RSFC features using ComBat-GAM (i.e., NeuroHarmonize) (41).

### Psychopathology features

Consistent with our previous work (1), we evaluated four key dimensions of psychopathology: positive symptoms, negative symptoms, general psychopathology, and manic symptoms. Specifically, 7 positive symptoms (e.g., hallucinations, delusions), 7 negative symptoms (e.g., avolition, anhedonia), and 16 general psychopathology symptoms (e.g., tension, somatic concern) were assessed by the Positive and Negative Syndrome Scale (PANSS) (42). 11 manic symptoms were evaluated by the Young Mania Rating Scale (YMRS) (43). This approach yielded a comprehensive set of 41 symptom-level features, providing a detailed psychopathological profile for each participant (Table S2). Additionally, ANCOVA was performed in Python to determine the differences of these clinical scores across diagnostic groups, while controlling for covariates including sex and age (Table S2; Figure S1).

### PLS analysis

We employed PLS correlation analysis to identify LCs maximizing covariation between functional connectivity and clinical features (Table S1-2). Prior to the PLS analysis, sex, age, and quadratic age were regressed out using Ordinary Least Squares regression. The PLS analysis methodology was consistent with our previous work (1). Briefly, this approach decomposes the brain and clinical data matrices into independent LCs characterized by maximum covariance, with each LC assigned a singular value representing the explained covariance between RSFC and symptoms (Supplemental Methods S8).

Significant LCs were determined through permutation testing (5,000 iterations) with false discovery rate (FDR) correction (*q*<0.05) (1). We calculated composite scores by projecting individual data onto these LCs, with loadings computed as Pearson’s correlations between composite scores and original features. Post-hoc analyses examined demographic and clinical factors potentially influencing LC composite scores (Supplemental Methods S9-10; Table S4).

We conducted a supplementary analysis comparing patients’ RSFC composite scores with healthy controls (Supplemental Methods S11) to determine whether identified connectivity patterns represented abnormal functional connectivity or within-normal-range variability.

For interpretation, we aggregated parcel-level loadings into 18 canonical networks, including 17 cortical networks from the Schaefer et al. (2018) atlas (Figure S2A) (39) and one subcortical network from the Tian et al. (2020) atlas (Figure S2B) (40). For each pair of networks, we averaged the parcel-level loadings for all connections within and between networks, producing an 18×18 network-level loading matrix representing mean contributions to the LC. This network-level aggregation facilitated interpretation of the multivariate patterns at a macroscale level (21).

### Associating Latent Symptom Clusters with Behavioral Measures

To further validate the clinical composite score of the latent component (LC) detected through PLS analysis, we examined its association with seven self-reported behavioral measures related to relevant traits and psychosocial stressors measured using the NIH Toolbox Emotion Battery (44). Specifically, we selected 1) Anger Survey, with subdomains of Physical Aggression, Affect, Hostility, and Negative Affect Summary scores (45); 2) Perceived Stress Scale (46); 3) Peer Rejection and Perceived Rejection (47); and 4) Psychological Well-being (48). In total, 7 measures including subdomains from these 4 surveys were included to assess whether they strengthen our understanding of the clinical features captured by the LC. Correlations between the clinical composite score of the detected LC and the 7 behavioral measures were calculated, with significance determined using Benjamini-Hochberg FDR correction (*q*<0.05).

## Results

### PLS Latent Component

PLS correlation was applied to detect the multivariate relationship between whole-brain RSFC and a comprehensive set of 41 behavioral measures in 124 patients across diagnostic categories. Notably, one single significant LC (*q*<0.001) captured 41.6% of the covariance between RSFC strength and clinical symptoms (Figure 1A). The RSFC and clinical composite scores of the detected LC were significantly correlated with each other (r=0.43, *p*<0.001; Figure 1B). No significant differences in RSFC strength or clinical composite scores were observed across diagnostic groups (i.e., schizophrenia, schizoaffective disorder, and psychotic mood disorders) after FDR correction (*q*>0.05) (Figure 1C-D), suggesting that the identified LC represents a transdiagnostic pattern of brain-behavior associations.

**Figure 1.**
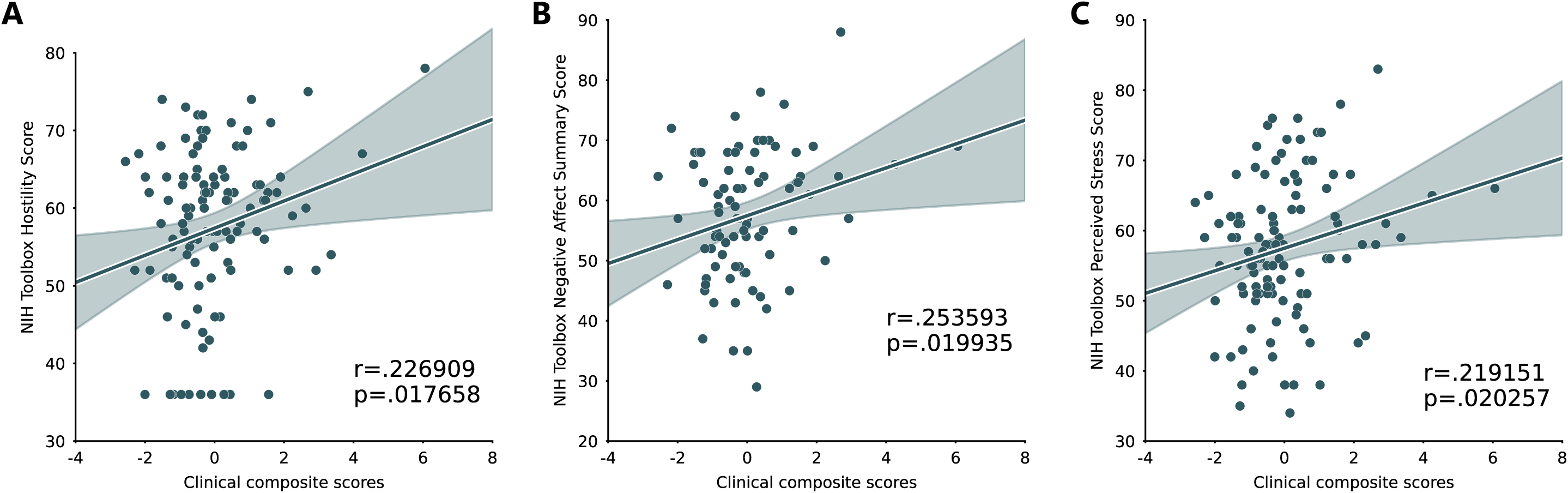
Primary Latent Component (LC) identified by PLS. **(A).** One single significant LC (*p*<.001) captures 41.6% of the covariance between RSFC variables and clinical symptoms. **(B).** The RSFC and clinical composite scores of the detected LC were significantly correlated with each other (r=0.43, *p*<.001). **(C).** Group differences in RSFC composite scores. No significant differences in RSFC composite scores were observed across diagnostic groups after FDR correction (*q*>0.05). **(D).** Group differences in clinical composite scores. No significant differences in clinical composite scores were observed across diagnostic groups after FDR correction (*q*>0.05).

### RSFC Loadings

Greater RSFC composite scores were characterized by a distributed pattern of between-network connections across the brain (Figure 2A-D). The strongest positive loadings were observed between the somatomotor networks and default mode networks (r=0.33-0.37), and between the visual network and default mode network (r=0.34-0.36). The subcortical regions also showed significant positive loadings in their connections with the salience and ventral attention network (r=0.33, SE=0.13), somatomotor networks (r=0.32-0.33), control B network (r=0.32, SE=0.12), and default mode networks (r=0.31-0.32). Specifically, among subcortical regions, the amygdala (r=0.41, SE=0.14), nucleus accumbens (r=0.38, SE=0.13), posterior thalamus (r=0.36, SE=0.12), and putamen (r=0.35, SE=0.12) demonstrated higher RSFC with sensory networks (visual and somatomotor) and default mode networks. Notably, the between-network RSFC loadings were generally stronger than within-network loadings, suggesting this LC primarily captures alterations in network integration rather than segregation. This loading pattern indicates that connections whose strength increases in correspondence with the LC composite score—i.e., greater RSFC strength associated with more pronounced clinical features—are predominantly between the default mode network, somatomotor networks, subcortical regions, and the control B network.

**Figure 2.**
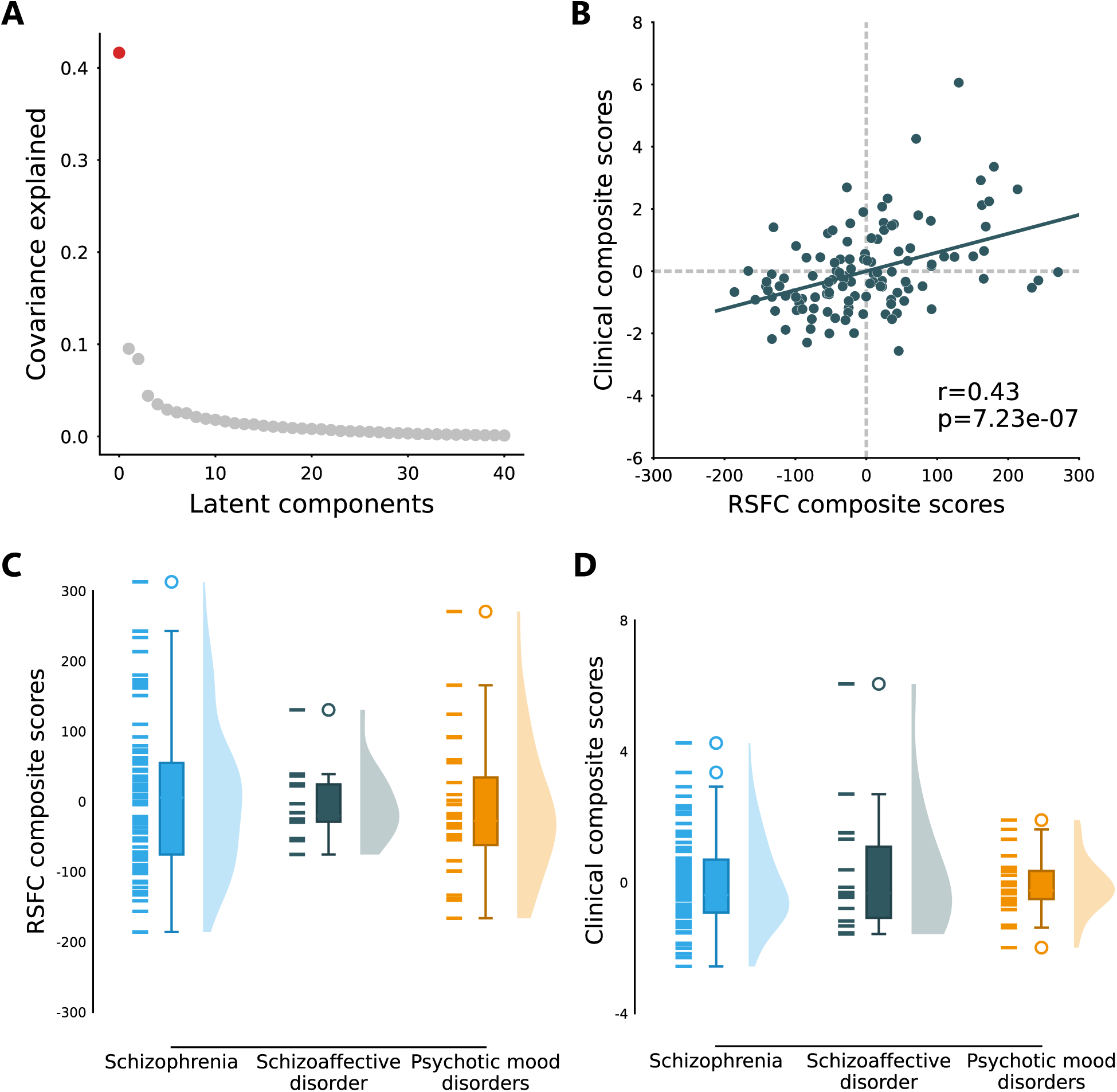
Profile of RSFC features that load onto the RSFC composite score of the detected LC. **(A)** Unthresholded correlations between participants’ RSFC features and their RSFC composite scores at the parcel level. Red (or blue) color indicates that greater RSFC is positively (or negatively) associated with LC. **(B)** Unthresholded correlations between participants’ RSFC data and their RSFC composite scores averaged within and between networks. **(C)** Thresholded significant RSFC loadings of the LC averaged by networks. Strong loadings with an effect size greater than 1 standard deviation were shown in the bar chart. Error bars indicate bootstrapped standard deviations. **(D)** Circle plot depicting significant within- and between-network RSFC loadings. Line color represents the magnitude of loading coefficients. Red (or blue) color indicates that greater RSFC is positively (or negatively) associated with LC.

To contextualize our findings within normative brain function to understand whether the RSFC strength composite pattern as related to this specific set of symptoms reflected variability within the normal range or represented higher than normal functional connectivity, we compared patients’ RSFC composite scores with those derived from the control group using the same LC weights. No significant differences were observed between patients and controls in RSFC composite scores (*p*>0.05) (Figure S3). This indicates that the functional coupling patterns we identified reflect connectivity variations associated with specific symptom expression within patients, rather than representing general hyperconnectivity relative to typical brain function.

### Clinical Loadings

In the PLS correlation analyses, clinical symptoms with significant loadings to the psychopathological pattern of the LC revealed a distinct profile characterized by cognitive rigidity and arousal dysregulation. Stereotyped thinking (r=0.19, SE=0.16), somatic concern (r=0.14, SE=0.13), anxiety (r=0.19, SE=0.16), and disruptive/aggressive behavior (r=0.16, SE=0.16) showed positive loadings, while hallucinatory behavior loaded negatively (r=-0.16, SE=0.15) (Figure 3A). This pattern suggests the RSFC signature was particularly associated with symptoms reflecting an inflexible cognitive style and heightened physiological tension, rather than perceptual disturbances. The positive loadings spanned multiple traditional symptom domains, combining negative symptoms (stereotyped thinking), general psychopathology (somatic concerns, anxiety), and behavioral reactivity (disruptive-aggressive behavior; Figure 3B; Table S3). The inverse relationship with hallucinatory behavior further distinguishes this symptom profile, suggesting it may capture a distinct profile characterized more by cognitive-somatic manifestations than by positive symptoms.

**Figure 3.**
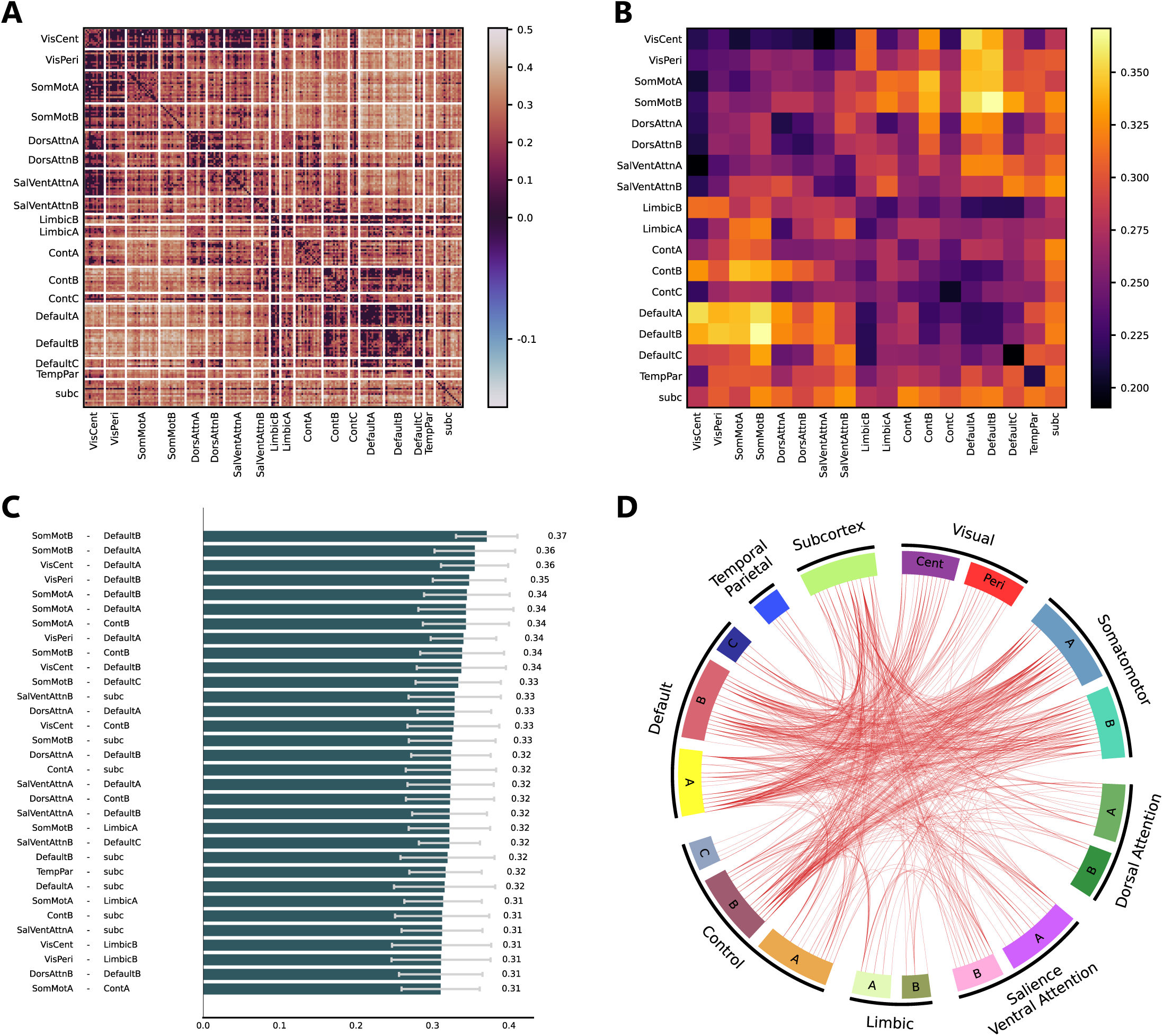
Profile of clinical features that load onto the clinical composite score of the detected LC. **(A)** Significant clinical loadings of the LC (*q*<0.05). Error bars indicate bootstrapped standard errors. **(B)** Unthresholded clinical loadings of the LC. Black, blue, green, and orange colors indicate the symptom dimensions of positive, negative, general, and mania symptoms, respectively. Significant loadings were outlined in red.

In contrast to the clinical composite scores identified by PLS, comparisons of raw clinical scores using ANCOVA revealed significant diagnostic differences in clinical scales (Table S2; Figure S1). PANSS positive symptoms differed across schizophrenia, schizoaffective disorder, and psychotic mood disorder (F(2,119)=10.19, *p*<0.001); PANSS negative symptoms were highest in schizophrenia versus schizoaffective and psychotic mood disorder (F(2,119)=6.05, *p*<0.01); and PANSS general psychopathology scores also differed (schizoaffective > schizophrenia > bipolar) (F(2,119)=4.35, *p*=0.02). No group difference was found in YMRS score (*p*>0.05).

### Correlation between clinical composite scores and emotion measures

To evaluate the clinical significance of the PLS-derived latent component (LC) and provide external validation for its symptom profile—characterized by cognitive inflexibility, hostility, and tension—we examined its relationship with standardized measures of emotional functioning from the NIH Toolbox Emotion Battery. These measures were selected for their relevance to assessing negative emotional states (e.g., hostility, perceived stress) and psychosocial stressors (e.g., peer rejection), which align with the LC’s clinical features. Significant positive correlations were found between the clinical composite score and three measures after FDR correction (Figure 4): hostility (r=0.23, *q*=0.047, Figure 4A), negative affect summary score (r=0.25, *q*=0.047, Figure 4B), and perceived stress (r=0.22, *q*=0.047, Figure 4C). Correlations with physical aggression, affect, peer rejection, and psychological well-being were not statistically significant (*q*>0.05). These significant associations with established measures of hostility, negative affect, and perceived stress provide external validation for the clinical relevance of the latent component, suggesting it captures meaningful aspects of psychopathology related to anger and negative emotional states and stress perception.

**Figure 4.**
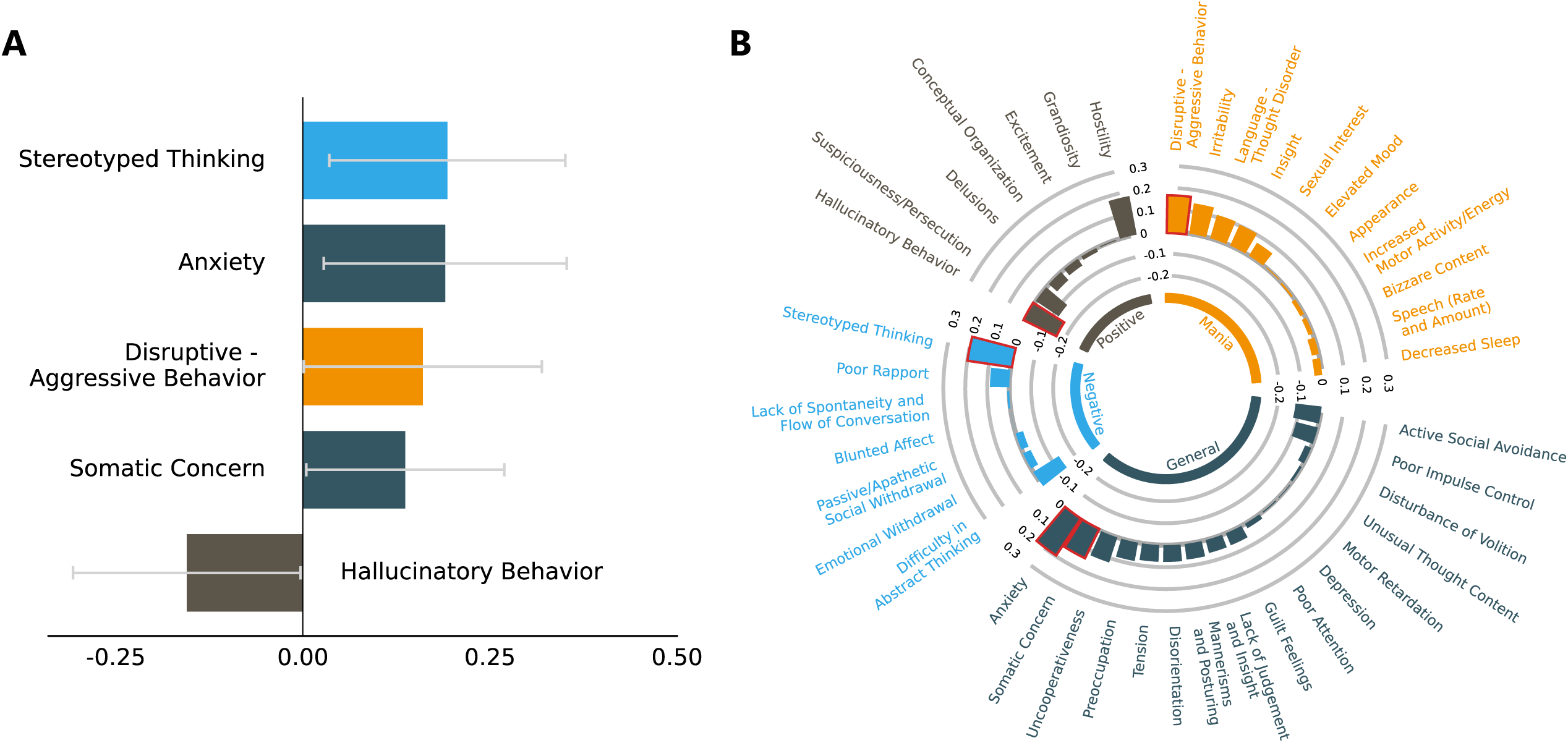
External validation of the PLS-derived latent component with standardized measures of emotional functioning. Individual participants’ clinical composite scores were correlated with NIH Toolbox Emotion Battery measures. Regression lines are plotted with 95% confidence interval shaded areas. **(A)** Significant positive correlation between clinical composite scores and hostility (r=0.23, *p*=0.018, *q*=0.047). **(B)** Significant positive correlation between the clinical composite score and negative affect summary score (r=0.25, *p*=0.20, *q*=0.047). **(C)** Significant positive correlation between clinical composite scores and perceived stress (r=0.22, *p*=0.20, *q*=0.047). All correlations survived FDR correction (*q*<0.05).

## Discussion

In this study, we employed a data-driven multivariate approach to investigate relationships between resting-state functional connectivity (RSFC) and psychopathology in early psychosis (EP). Our analyses revealed a single significant latent component that captured 41.8% of the covariance between RSFC strength and clinical features. This component was characterized by a distributed pattern of greater functional connectivity, particularly involving interactions between sensory-motor networks, default mode networks, and subcortical regions including the amygdala, nucleus accumbens, and thalamus. The clinical profile associated with this RSFC pattern was marked by symptoms that broadly reflect cognitive rigidity (49, 50) and arousal dysregulation (51–53), including stereotyped thinking, anxiety, and somatic concerns, rather than traditional positive symptoms like hallucinations or negative symptoms like social withdrawal, anhedonia, and avolition. Notably, this brain-behavior relationship was consistent across diagnostic categories and independent of potential confounds such as psychotropic medication and substance use. The clinical relevance of this latent component was further validated through its significant associations with standardized measures of emotional functioning, particularly hostility, negative affect, and perceived stress. These findings suggest that specific patterns of network-level functional connectivity may relate to distinct symptom dimensions that cut across conventional diagnostic boundaries in EP.

The distributed pattern of functional connectivity identified in our analyses highlights the complex interplay between multiple brain systems in EP. The somatomotor and visual networks, which showed significant positive connectivity with both default mode and subcortical regions, are fundamental for basic sensory processing and motor control (20, 21), potentially reflecting altered sensory integration and motor regulation that has been previously observed in psychosis (23, 54, 55). The default mode network, traditionally associated with self-referential processing and internal state monitoring (11, 56), demonstrated enhanced connectivity with both sensory and subcortical systems, suggesting disrupted integration between internal and external processing streams (4, 9). This mirrors longitudinal EP findings, which demonstrate that RSFC disturbances—such as heightened thalamus–sensorimotor coupling and weakened frontoparietal integration—undergo dynamic changes, often intensifying or shifting across networks during the first years following illness onset (5, 23). The involvement of subcortical regions, particularly the amygdala, nucleus accumbens, and thalamus, is noteworthy given the crucial roles of these structures in emotional processing, reward, and sensory gating (6, 57). Although mean RSFC composite scores did not differ from healthy controls, RSFC strength within these circuits were positively associated with symptom severity. This implies that variation within the normative range of functional coupling—rather than absolute hyperconnectivity—may index dysregulated arousal and emotional processing systems underlying cognitive rigidity and tension. These networks have been previously implicated in various aspects of psychosis, including cognitive control deficits (58), emotional processing alterations (59), and disturbances in self-referential processing (60). Our findings extend this work by demonstrating how these same neural circuits, known to support various cognitive and affective functions, may also underlie a distinct symptom profile characterized by cognitive inflexibility and arousal dysregulation.

Notably, our findings emphasized between-network rather than within-network functional connectivity patterns, particularly involving interactions between sensory-motor, default mode, and subcortical systems. Greater between-network RSFC strength in our analyses reflects stronger functional coupling between these systems in relation to specific symptom severity, rather than hyperconnectivity relative to typical brain function. This distinction is important, as our analyses examined functional connectivity patterns within the patient sample that correlate with symptom expression, not case-control differences. Previous studies have shown that altered between-network connectivity in psychosis may reflect disrupted network segregation (10) or compromised integration of information across brain systems (4). The predominance of between-network connections in our findings, particularly involving sensory-motor systems with both higher-order networks (default mode) and subcortical regions, suggests that symptoms of cognitive rigidity and arousal dysregulation may arise from altered coordination between basic sensory processing and higher-level cognitive control systems (21). This aligns with theories proposing that psychosis involves disrupted integration of external and internal information processing (9). The involvement of subcortical connections further suggests that this disrupted integration may be particularly relevant for emotional and arousal regulation (57). Importantly, these connectivity patterns represent relative differences in functional coupling strength across different brain systems within individuals, corresponding to symptom severity, rather than absolute measures of functional connectivity strength compared to a control population.

The symptom profile identified in our analyses presents a distinct clinical phenotype that diverges from traditional characterizations of psychosis symptomatology. Rather than emphasizing positive symptoms such as hallucinations and delusions, or negative symptoms such as diminished expression, this profile encompasses a constellation of features centered on cognitive rigidity, arousal dysregulation, and somatic tension. This divergence aligns with our transdiagnostic approach, which examined different clinical presentations (which may or may not have included high levels of positive and negative symptoms) to find core clinical patterns based on shared associations with RSFC patterns. Notably, this symptom constellation has some parallels to trauma-related presentations, consistent with research indicating approximately 40% of first-episode psychosis patients report a history of significant traumatic exposure (61). This connection is further supported by the significant associations between our clinical composite scores and standardized measures of perceived stress, hostility, and negative affect—aligning with evidence that emotional dysregulation in psychosis is influenced by factors including current stressors and heightened arousal (62). These findings accord with broader consensus suggesting that trauma exposure is associated with elevated negative emotional states and stress sensitivity in clinical high-risk populations, which subsequently predict conversion to psychosis (63–65). The neurobiologically-grounded approach to phenotyping complements traditional diagnostic frameworks may indicate that stress regulation and cognitive flexibility are linked and may be important treatment target (66). While trauma assessment was not included in the HCP dataset, our findings highlight the clinical utility of identifying transdiagnostic symptom clusters that may respond to specific therapeutic approaches. Future research integrating direct assessment of trauma history alongside neuroimaging could further clarify these relationships, demonstrating that attention to these specific symptom dimensions may be valuable for understanding and treating EP, even in the absence of overt positive symptoms.

Several methodological strengths enhance the validity and reliability of our findings. First, our use of Partial Least Squares correlation provided key advantages over traditional univariate approaches by enabling simultaneous examination of multiple brain-behavior relationships while accounting for their inherent correlations (21, 29). This multivariate framework was particularly valuable for detecting subtle but distributed patterns of functional connectivity that may not be apparent when examining individual connections in isolation. Second, rather than relying solely on diagnostic categories or clinical symptoms, we validated our findings through independent behavioral measures from the NIH Toolbox, providing external confirmation of the clinical relevance of our identified component. The significant correlations between our clinical composite scores and standardized measures of emotional functioning strengthen the interpretation of our results beyond the primary clinical variables. Third, our analyses carefully controlled for potential confounding factors including age, sex, education, head motion, substance use, and medication effects, with results remaining robust after accounting for these variables. Specific to the RSFC analyses, we demonstrated that our findings were stable across different preprocessing approaches, including analyses with and without global signal regression, supporting the robustness of the identified brain-behavior relationships (Figure S4-6). This comprehensive methodological approach enhances confidence in the reliability and generalizability of our results (12).

### Limitations and conclusions

Several limitations of the current study should be noted. First, while the sample size was adequate for multivariate analyses, replication in independent cohorts will be crucial for establishing the generalizability of our findings across different early psychosis populations. Second, the cross-sectional nature of the data precludes conclusions about the temporal dynamics of these brain-behavior relationships - longitudinal studies will be crucial for understanding how these patterns may evolve over the course of illness. Third, the lack of trauma history data in the sample, while not uncommon in psychosis research, limits our ability to directly test hypotheses about trauma’s role in the observed symptom profile and RSFC patterns. Despite these limitations, our findings provide a robust foundation for future investigations into the relationship between functional brain network organization and symptom expression in EP.

In conclusion, our study highlights how rs-fMRI—a method uniquely capable of capturing dynamic, state-dependent functional connectivity patterns rather than static deficits—can reveal clinically meaningful brain-behavior relationships in EP. We identified a specific transdiagnostic symptom cluster characterized by cognitive rigidity, arousal dysregulation, and somatic concerns, associated with a distinct functional connectivity signature involving sensory-motor, default mode, and subcortical systems. This approach complements traditional diagnostic frameworks by detecting meaningful clinical variability and patterns that might otherwise be overlooked. Our findings should be viewed as a snapshot of an evolving neurobiological process specific to the EP phase, where intervention might be most impactful. Future longitudinal research examining how these brain-behavior relationships evolve over time and respond to targeted interventions will be essential for translating these insights into improved care that address this previously underemphasized symptom dimension.

## Author contributions

Author Contributions: Haley Wang had full access to all of the data in the study and takes responsibility for the integrity of the data and the accuracy of the data analysis.

Study concept and design: HRW, KHK. Acquisition, analysis, or interpretation of data: HRW, ZQL, CS, JSN, CEB, LQU, KHK. Drafting of the manuscript: HRW, KHK. Critical revision of the manuscript for important intellectual content: HRW, ZQL, CEB, LQU, BM, KHK. Statistical analysis: HRW, ZQL, KHK. Obtained funding: KHK. Administrative, technical, or material support: ZQL, CS, JSN, CEB, LQU, BM, KHK. Study supervision: HRW, KHK.

## Acknowledgement

Research using Human Connectome Project for Early Psychosis (HCP-EP) data reported in this publication was supported by the National Institute of Mental Health of the National Institutes of Health under Award Number U01MH109977. The HCP-EP 1.1 Release data used in this report came from DOI: 10.15154/1522899. This work used computational and storage services associated with the Hoffman2 Shared Cluster provided by UCLA Institute for Digital Research and Education’s Research Technology Group.

## Data Sharing Statement

All data and code used to perform the analyses can be found at https://github.com/haleyrwang/CCNL_RSFC_PLS. The HCP-EP dataset is available at https://db.humanconnectome.org/. The volumetric PET images and receptor maps are available via neuromaps (https://github.com/netneurolab/neuromaps) (61).

