## Supplemental material for "Multivariate Resting-State Functional Connectivity Features Linked to Transdiagnostic Psychopathology in Early Psychosis"

***Supplemental Information***

**Supplemental Methods**

**Methods S1.** Participant details

**Methods S2.** Differences in symptom severity across diagnostic groups

**Methods S3.** Categorization and dosage of medication

**Methods S4.** Substance use

**Methods S5.** MRI data acquisition

**Methods S6.** Resting state imaging data analysis

**Methods S7.** Global signal regression (GSR) for RSFC features

**Methods S8.** Partial least squares (PLS) correlation analysis details

**Methods S9.** Sensitivity analysis and robustness control

**Methods S10.** Post-hoc analyses on the associations between composite scores and confounds

**Methods S11.** Comparison of RSFC composite scores between patients and controls

**Supplemental Results**

**Table S1.** All clinical measures used in the PLS analyses

**Table S2.** Group scores on all clinical measures included in the PLS analyses

**Figure S1.** Symptom severity variations by diagnostic category

**Table S3.** Associations between clinical features and clinical composite scores

**Figure S2.** Brain parcellation

**Figure S3.** Comparison of RSFC composite scores between patients and controls

**Figure S4*.*** Primary Latent Component (LC) identified by PLS with GSR

**Figure S5.** Profile of RSFC features that load onto the RSFC composite score of the detected LC with GSR

**Figure S6.** Profile of clinical features that load onto the clinical composite score of the detected LC with GSR

**Table S4.** Associations between RSFC (or clinical) composite scores and potential confounds

**Supplemental References**

This supplementary material has been provided by the authors to give readers additional information about their work.

***Methods S1.*** *Participant details*

This investigation utilized data from the Human Connectome Project for Early Psychosis (HCP-EP, release 1.1), comprising 182 participants (124 individuals with psychosis and 58 demographically matched unaffected controls) between ages 16-35 at enrollment. The clinical sample included participants meeting diagnostic criteria according to the Diagnostic and Statistical Manual of Mental Disorders (DSM-5) for either primary psychotic disorders (schizophrenia, schizophreniform, schizoaffective, psychosis NOS, delusional disorder, or brief psychotic disorder) or mood disorders with psychotic features (major depression with psychosis or bipolar disorder with psychosis). A key inclusion criterion was psychosis onset within three years prior to study participation. Exclusion criteria encompassed intellectual disability (IQ below 70), history of significant neurological conditions or traumatic brain injury, and contraindications for MRI assessment. Three participants were excluded from neuroimaging analyses due to absent T1-weighted structural scans, and an additional three were excluded from clinical analyses owing to incomplete psychiatric symptom assessments. All assessments were conducted by the HCP research team following protocols detailed in the HCP-EP_Release_1.1_Manual. Symptom evaluation in the psychosis group was performed using the Positive and Negative Syndrome Scale (PANSS)(1) and Young Mania Rating Scale (YMRS)(2).

***Methods S2.*** *Differences in symptom severity across diagnostic groups*

We analyzed variations in clinical presentations across diagnostic categories included in our PLS analysis. Utilizing clinical data from the HCP-EP cohort, we compared symptom profiles using total scores from the PANSS positive, negative, and general symptom subscales, as well as YMRS scores (Table S1). Statistical analyses were conducted in Python, implementing ANOVA to evaluate significant clinical differences between diagnostic groups while adjusting for demographic covariates including participant sex and age (Table S2; Figure S1).

***Methods S3.*** *Categorization and dosage of medication*

Antipsychotic medication dosages were standardized by conversion to chlorpromazine equivalent values (3). We examined potential relationships between these standardized medication loads and both resting-state functional connectivity (RSFC) and clinical composite scores using Pearson correlation analyses (Table 2). The study did not account for other pharmacological treatments such as mood stabilizers, antidepressants, anxiolytics/sedatives/hypnotics, or stimulants, as comprehensive medication information was not available in the public datasets utilized.

***Methods S4.*** *Substance use*

HCP-EP participants underwent substance use screening on the day of their scanning session. Individuals with substance use disorders included in the study presented either mild symptoms or were in remission. For our analyses, substance use was specifically defined as cannabis consumption within the preceding 30 days, a focus selected due to the elevated prevalence of cannabis use among individuals with psychotic disorders. Cannabis use status was categorized dichotomously as either present or absent, without further quantification of usage frequency. We evaluated potential associations between cannabis use status and both RSFC and clinical composite scores using independent two-sample t-tests (Table 2).

***Methods S5.*** *MRI data acquisition*

Imaging data of the HCP-EP study were collected on three Siemens MAGNETOM Prisma 3T scanners across 4 sites. Scanning sessions included a collection of T1-weighted (multi-echo 3D MPRAGE, TR/TE/TI 2400/2.22/1000ms, 8° flip angle, FOV 256mm, 0.8mm isotropic resolution, 2x GRAPPA acceleration, 208 sagittal slices) and T2-weighted (3D variable-flip-angle turbo-spin-echo (TSE) sequence SPACE, TR/TE 3200/563ms, FOV 256mm, 0.8mm isotropic, 2x GRAPPA, turbo factor 314ms, 208 sagittal slices) structural scans. Additionally, spin echo field maps were collected 4 times with twice for each AP and PA phase encoding (TR/TE 8000/66ms, FoV 208mm, 90° flip angle, 2mm isotropic resolution). Resting state functional MRI data were acquired using a multiband echo-planar imaging sequence (TR/TE 800/37ms, 52° flip angle, FOV 208mm, 2mm isotropic resolution, multiband acceleration factor 8). Four runs were collected, with two runs using anterior-to-posterior and two runs using posterior-to-anterior phase encoding directions. Each run had an acquisition time of 5:47 minutes.

***Methods S6.*** *Resting-state imaging data analysis*

Raw structural and resting-state functional MRI (rs-fMRI) data in NIfTI format were acquired from HCP-EP study. Preprocessing and quality control for HCP-EP were performed using HCP Pipelines(4) implemented through Quantitative Neuroimaging Environment & Toolbox (QuNex)(5), which is compatible with multi-band and single-band fMRI. The fMRI time series underwent additional processing steps including bandpass filtering; motion scrubbing for frames that exceeded either framewise displacement or signal change thresholds; and spatial smoothing. Global signal regression (GSR) was not performed for the main analyses. Three patients were excluded due to missing T1w images. Any scans where more than 50% of frames were flagged for motion were removed from analysis. Additionally, ComBat-GAM (i.e., NeuroHarmonize) (6) was employed to harmonize RSFC data to adjust for scanner difference in the sample.

***Methods S7.*** *Global signal regression (GSR) for RSFC features*

While global signal regression (GSR) was not performed in our main analyses due to evidence that global signal correlates with key clinical and behavioral measures in psychosis (7, 8), we conducted supplementary analyses with GSR to test the robustness of our findings. For these supplementary analyses, the same preprocessing pipeline described in Methods S6 was applied, with the addition of regression of mean gray matter signal. This approach allowed us to address the ongoing debate regarding GSR in neuroimaging studies while preserving potentially important disease-related variance in our primary results. The supplementary GSR analyses (Figure S4-6) followed identical subsequent analytical steps as the main analyses to enable direct comparison of results with and without GSR.

***Methods S8.*** *Partial least squares analysis*

We employed partial least squares (PLS) correlation to identify linear relationships between 23,328 brain RSFC features and 41 psychopathology measures using the Pyls 0.1.7 Python package. This unsupervised approach decomposes the cross-product matrix via singular value decomposition, yielding RSFC loadings, clinical loadings, and singular values representing relationship strength. Prior to analysis, we controlled for age, quadratic age, and sex through linear regression and z-transformed the residuals.

Feature contribution was assessed using squared-loadings with z-scores above 2 indicating significant impact. Statistical significance was determined through permutation testing (5,000 repetitions) of singular values, while feature stability was evaluated using bootstrap resampling with 95% confidence intervals. Split-half resampling (10,000 iterations) was conducted to examine loading reproducibility. For comprehensive analytical details, see Wang et al. (9).

***Methods S9.*** *Robustness control*

To validate our findings' robustness, we implemented multiple verification strategies. Bootstrap resampling techniques were applied to all loadings, strengthening their stability and confirming result reliability. We examined potential multivariate outliers within the PLS outcome data using Mahalanobis distance calculations—an approach well-suited for multidimensional data analysis. This assessment revealed no data points exceeding our predetermined threshold of 2 standard deviations, eliminating the need for outlier removal. To address concerns about potential artifacts from data harmonization, we conducted correlation analyses comparing PLS outcomes before and after neuroHarmonize processing. These analyses demonstrated strongly correlated RSFC composite scores between models (p<.001), confirming that our harmonization approach preserved the underlying signal patterns.

***Methods S10.*** *Post-hoc analyses on the associations between composite scores and confounds*

We evaluated potential associations between the derived RSFC and clinical composite scores from the PSL-derived latent components and several variables of interest. These included demographic factors (sex, age, quadratic age), clinical characteristics (diagnostic categories, illness duration), methodological variables (data collection site, head motion parameters), and clinical confounds (substance use history, antipsychotic medication dosage). For continuous variables, we calculated Pearson correlation coefficients, while binary variables were assessed using independent t-tests (Table 2). These analyses incorporated all participants from the PLS analysis cohort, with false discovery rate (FDR) correction (q<.05) applied across all statistical comparisons to control for multiple testing.

***Methods S11.*** *Comparison of RSFC composite scores between patients and controls*

To determine whether the RSFC patterns identified in our PLS analysis represented abnormal functional connectivity relative to typical brain function, we compared patients' RSFC composite scores with those of demographically matched healthy controls from the HCP-EP 1.1 dataset. Using the methodology described by Wang et al. (2024) (9), we applied the weights derived from our patient-only PLS analysis to the RSFC data from healthy controls.

Specifically, we extracted the weights of each RSFC feature from the significant latent component (LC) identified in our patient PLS analysis. We then multiplied these PLS-derived weights by the raw, individual-level RSFC metrics in the control subjects to compute projected RSFC composite scores for each healthy control participant. This approach allowed us to quantify the expression of the patient-derived connectivity pattern in individuals without psychosis.

Subsequently, we conducted independent t-tests to assess group differences between patient and control RSFC composite scores. This analysis enabled us to determine whether the functional connectivity patterns associated with symptom severity in patients represented altered connectivity outside the normative range, or alternatively, whether they reflected symptom-related variability occurring within typical connectivity parameters.

**Table S1.** All clinical measures used in the PLS analyses. This table presents the clinical assessments available for study participants, indicating both complete measures included in the primary PLS analysis and measures with incomplete data (denoted by *) that required median imputation. The table categorizes each item from the original clinical scales according to the psychopathology dimensions defined in the study methodology.

| **Scale** | **Subscale/Domain**  (Defined in scale) | **Symptom** | **Dimension**  (Defined in study) | **Missing subjects** |
| --- | --- | --- | --- | --- |
| PANSS | Positive | Delusions | Positive | NA |
|  |  | Conceptual Organization | Positive | NA |
|  |  | Hallucinatory Behavior | Positive | NA |
|  |  | Excitement | Positive | NA |
|  |  | Grandiosity | Positive | NA |
|  |  | Suspiciousness/Persecution | Positive | NA |
|  |  | Hostility | Positive | NA |
|  | Negative | Blunted Affect | Negative | NA |
|  |  | Emotional Withdrawal | Negative | *(2) |
|  |  | Poor Rapport | Negative | *(1) |
|  |  | Passive/Apathetic Social Withdrawal | Negative | *(1) |
|  |  | Difficulty in Abstract Thinking | Negative | *(2) |
|  |  | Lack of Spontaneity and Flow of Conversation | Negative | *(1) |
|  |  | Stereotyped Thinking | Negative | *(1) |
|  | General | Somatic Concern | General | NA |
|  |  | Anxiety | General | *(1) |
|  |  | Guilt Feelings | General | *(1) |
|  |  | Tension | General | NA |
|  |  | Mannerisms and Posturing | General | *(1) |
|  |  | Depression | General | *(1) |
|  |  | Motor Retardation | General | NA |
|  |  | Uncooperativeness | General | NA |
|  |  | Unusual Thought Content | General | *(1) |
|  |  | Disorientation | General | *(1) |
|  |  | Poor Attention | General | *(1) |
|  |  | Lack of Judgement and Insight | General | *(1) |
|  |  | Disturbance of Volition | General | *(1) |
|  |  | Poor Impulse Control | General | NA |
|  |  | Preoccupation | General | *(1) |
|  |  | Active Social Avoidance | General | *(2) |
| YMRS | Mania | Elevated Mood | Mania | NA |
|  |  | Increased Motor Activity-Energy | Mania | NA |
|  |  | Irritability | Mania | NA |
|  |  | Language-Thought Disorder | Mania | NA |
|  |  | Content | Mania | NA |
|  |  | Disruptive-Aggressive Behavior | Mania | NA |
|  |  | Appearance | Mania | NA |
|  |  | Insight | Mania | NA |
|  |  | Increased Speech (Rate and Amount) | Mania | NA |
|  |  | Increased Sexual Interest | Mania | NA |
|  |  | Sleep | Mania | NA |

**Table S2.** Diagnostic differences in symptom severity. Mean (SD) values of the summarized clinical scores are presented for each primary diagnostic group. Groups were compared using ANCOVA with age and sex included as covariates. P-values that remained significant after FDR correction (q<.05) are indicated with *.

| Scale / Subscale | | SZ | SZAD | BD | F | p value |
| --- | --- | --- | --- | --- | --- | --- |
| PANSS | Positive | 11.54 (3.77) | 14 (4.9) | 8.89 (2.47) | 10.19 | <.001* |
|  | Negative | 15.3 (5.77) | 12.71 (3.27) | 10.96 (3.72) | 6.05 | <.01* |
|  | General | 25.43 (4.94) | 25.64 (6.25) | 22 (4.87) | 4.35 | 0.02* |
| YMRS | Mania | 4.5 (4.52) | 6.79 (8.42) | 4.98 (5.39) | 1.07 | 0.35 |

*Abbreviations:* *SZ=schizophrenia spectrum disorders, including schizophrenia, schizophreniform, psychosis NOS, delusional disorder, or brief psychotic disorder. SZAD=schizoaffective disorder; BP= bipolar disorders with psychotic features or major depressive disorder with psychotic features.*

**Figure S1.** Symptom severity variations by diagnostic category. Within the HCP-EP cohort, analyses revealed significant severity differences across diagnostic groups (schizophrenia, schizoaffective disorder, and psychotic mood disorders) for positive, negative, and general symptom domains after FDR correction. No significant between-group differences were detected for mania symptoms (statistical details available in Table S2, which presents ANCOVA results).

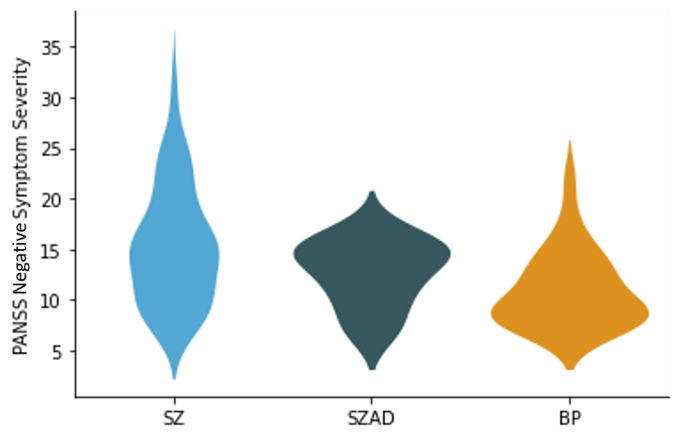

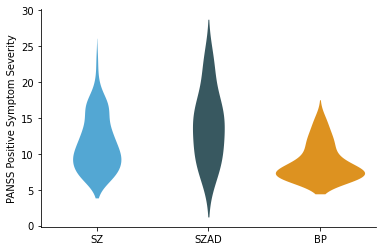
**A	B**

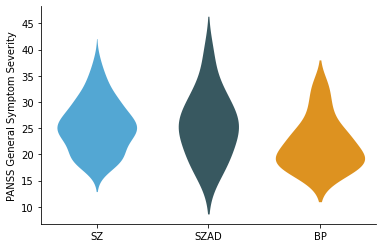

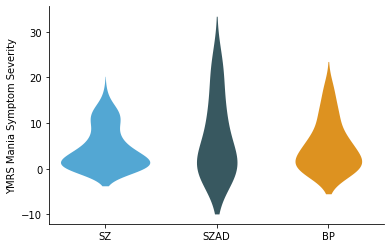
**C	D**

**Figure S2.** Brain parcellation methodology. **(A).** Cortical regions based on Schaefer's 200-parcel atlas. These parcels are organized into 17 resting-state networks (10). Note that the 200 cortical parcellations are not symmetric between hemispheres. **(B).** Subcortical structures comprising 16 regions (11). These combined parcellations yielded 216 × 216 RSFC matrices for each participant. Due to the symmetrical nature of RSFC matrices, analyses utilized only the upper triangular portions (though complete matrices are displayed for visualization purposes).

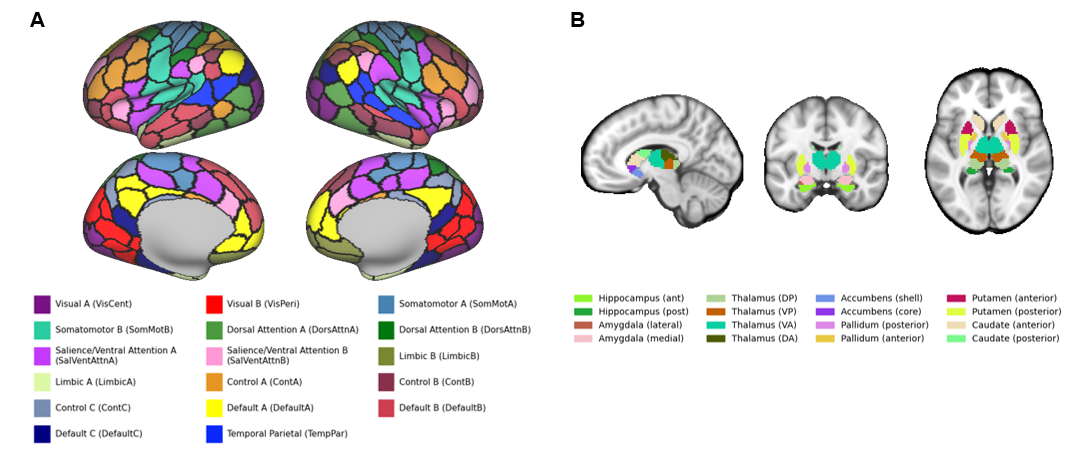

**Table S3.** Correlation analysis between clinical measures and clinical composite scores. This table presents Pearson's correlation coefficients demonstrating each clinical measure's contribution to the identified latent component, accompanied by bootstrap-derived standard errors (SE). Asterisks (*) indicate correlations with statistically significant bootstrapped Z scores (p<.05).

| **Scale** | **Dimension** (Defined in study) | **Feature** | **Corr (SE)** |
| --- | --- | --- | --- |
| PANSS | Positive | Delusions | -0.05 (0.17) |
|  |  | Conceptual Organization | -0.04 (0.17) |
|  |  | Hallucinatory Behavior | -0.16 (0.15)* |
|  |  | Excitement | -0.02 (0.16) |
|  |  | Grandiosity | -0.00 (0.16) |
|  |  | Suspiciousness/Persecution | -0.12 (0.16) |
|  |  | Hostility | 0.18 (0.18) |
|  | Negative | Blunted Affect | 0.00 (0.19) |
|  |  | Emotional Withdrawal | -0.04 (0.20) |
|  |  | Poor Rapport | 0.09 (0.21) |
|  |  | Passive/Apathetic Social Withdrawal | -0.03 (0.16) |
|  |  | Difficulty in Abstract Thinking | -0.14 (0.17) |
|  |  | Lack of Spontaneity and Flow of Conversation | 0.01 (0.18) |
|  |  | Stereotyped Thinking | 0.19 (0.16)* |
|  | General | Somatic Concern | 0.14 (0.13)* |
|  |  | Anxiety | 0.19 (0.16)* |
|  |  | Guilt Feelings | 0.06 (0.15) |
|  |  | Tension | 0.08 (0.17) |
|  |  | Mannerisms and Posturing | 0.07 (0.19) |
|  |  | Depression | 0.00 (0.15) |
|  |  | Motor Retardation | 0.00 (0.17) |
|  |  | Uncooperativeness | 0.12 (0.23) |
|  |  | Unusual Thought Content | -0.01 (0.16) |
|  |  | Disorientation | 0.08 (0.16) |
|  |  | Poor Attention | 0.02 (0.15) |
|  |  | Lack of Judgement and Insight | 0.06 (0.18) |
|  |  | Disturbance of Volition | -0.03 (0.18) |
|  |  | Poor Impulse Control | -0.10 (0.12) |
|  |  | Preoccupation | 0.08 (0.23) |
|  |  | Active Social Avoidance | -0.12 (0.13) |
| YMRS | Mania | Elevated Mood | 0.00 (0.15) |
|  |  | Increased Motor Activity-Energy | -0.02 (0.14) |
|  |  | Increased Sexual Interest | 0.07 (0.19) |
|  |  | Decreased Sleep | -0.03 (0.15) |
|  |  | Irritability | 0.14 (0.17) |
|  |  | Increased Speech (Rate and Amount) | -0.03 (0.14) |
|  |  | Language-Thought Disorder | 0.11 (0.19) |
|  |  | Bizarre Content | -0.02 (0.16) |
|  |  | Disruptive-Aggressive Behavior | 0.16 (0.16)* |
|  |  | Appearance | -0.01 (0.15) |
|  |  | Insight | 0.10 (0.22) |

**Figure S3.** Comparison of RSFC composite scores between patients and controls. Indepentdent t-test was used to compare patients' RSFC composite scores to those derived from the control group using the same LC weights. No significant differences were observed between patients and controls in RSFC composite scores (*p*>0.05).

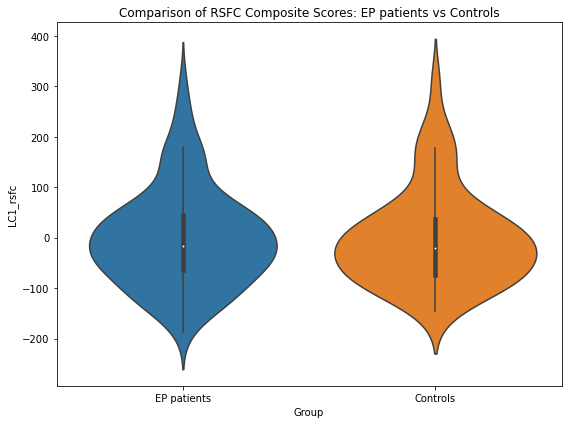

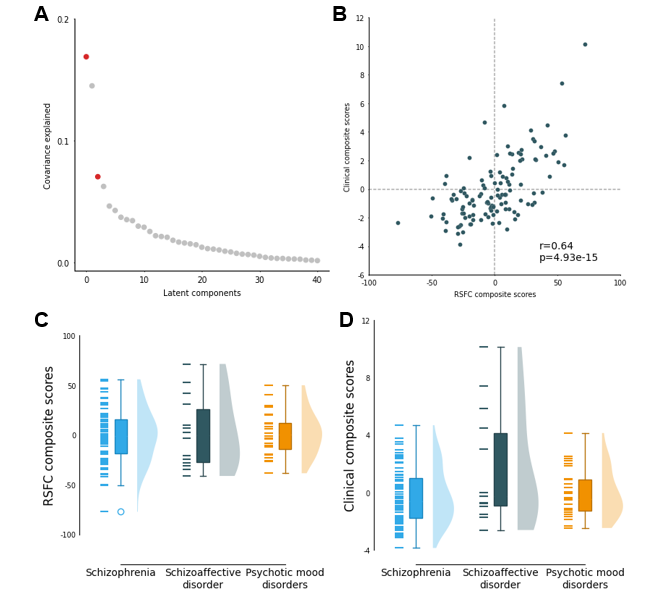
**Figure S4*.*** Primary Latent Component (LC) identified by PLS with GSR.

**(A).** Two significant LCs captures 16.9% (p=0.008) and 7% (p=0.027) of the covariance between RSFC variables and clinical symptoms, respectively. Due to the same percentage of covariance explained by LC2, we only show the results of loadings of LC1 which accounts for 16.9% of covariance between RSFC strength after GSR and clinical features. **(B).** The RSFC and clinical composite scores of LC1 were significantly correlated with each other (r=.64, *p*<.001). **(C).** Group differences in RSFC composite scores. No significant differences in RSFC composite scores were observed across diagnostic groups after FDR correction (*q*>0.05). **(D).** Group differences in clinical composite scores. No significant differences in clinical composite scores were observed across diagnostic groups after FDR correction (*q*>0.05).

**Figure S5.** Profile of RSFC features that load onto the RSFC composite score of LC1 with GSR.

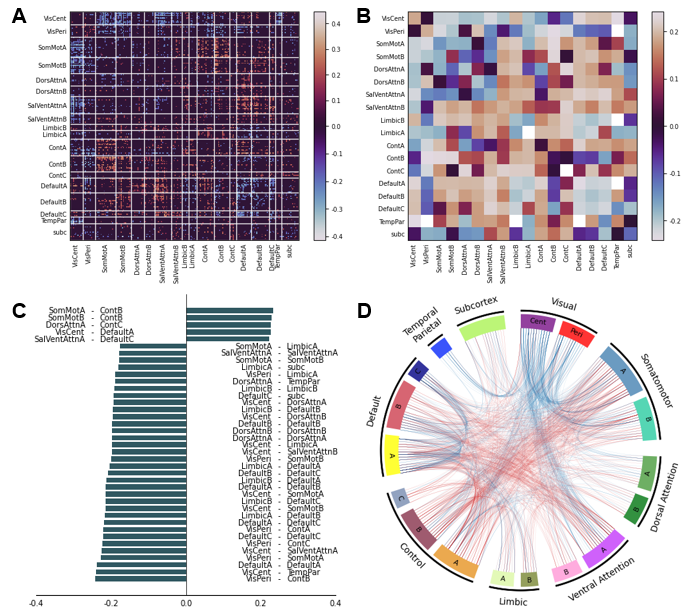

**(A)** Unthresholded correlations between participants' RSFC features and their RSFC composite scores at the parcel level. Red (or blue) color indicates that greater RSFC is positively (or negatively) associated with LC. **(B)** Unthresholded correlations between participants' RSFC data and their RSFC composite scores averaged within and between networks. **(C)** Thresholded significant RSFC loadings of the LC averaged by networks. Strong loadings with an effect size greater than 1.2 standard deviation were shown in the bar chart. Error bars indicate bootstrapped standard deviations. **(D)** Circle plot depicting significant within- and between-network RSFC loadings. Line color represents the magnitude of loading coefficients. Red (or blue) color indicates that greater RSFC is positively (or negatively) associated with LC.

**Figure S6.** Profile of clinical features that load onto the clinical composite score of LC1 with GSR.

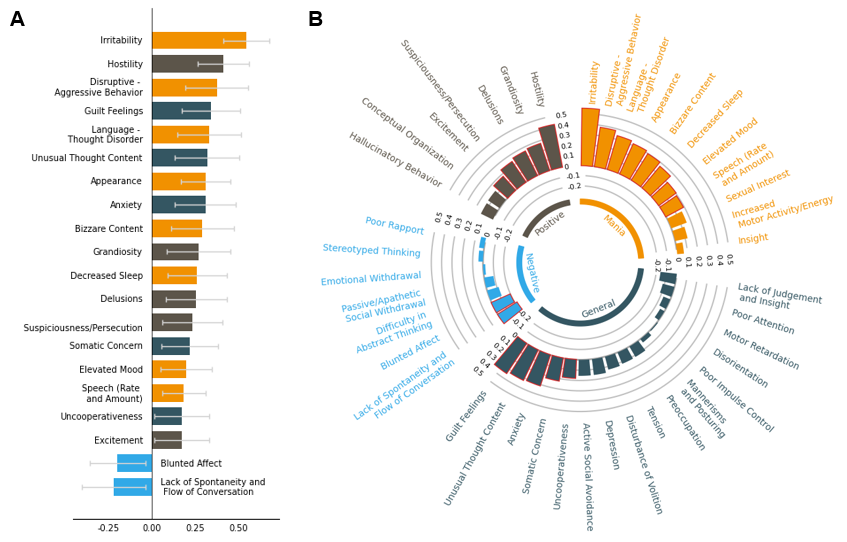

**(A)** Significant clinical loadings of the LC (*q*<0.05). Error bars indicate bootstrapped standard errors. **(B)** Unthresholded clinical loadings of the LC. Black, blue, green, and orange colors indicate the symptom dimensions of positive, negative, general, and mania symptoms, respectively. Significant loadings were outlined in red. Overall, the significant the LC identified in PLS analysis with GSR on RSFC features capture highly similar clinical loadings compared to the non-GSR PLS results reported in the main results.

**Table S4.** Associations between RSFC (or clinical) composite scores and potential confounds. We assessed associations between composite scores and potential confounds including age, sex, IQ, motion parameters, substance use, duration of psychosis, medication, and diagnosis. Both RSFC and clinical composite scores showed no significant correlations with confounds after FDR correction (q>0.05), indicating the independence of LC from confounds (Table S4).

|  | RSFC composite score | | Clinical composite score | |
| --- | --- | --- | --- | --- |
|  | *r/t* | *q* | *r/t* | *q* |
| Age | 0.00 | 1.00 | 0.00 | 1.00 |
| Sex | 0.00 | 1.00 | 0.00 | 1.00 |
| IQ | 0.00 | 1.00 | 0.00 | 1.00 |
| Site | 0.00 | 1.00 | 0.00 | 1.00 |
| Motion | 0.00 | 1.00 | 0.00 | 1.00 |
| Substance use | 0.48 | 1.00 | -0.72 | 1.00 |
| Duration of illness | 0.02 | 1.00 | 0.00 | 1.00 |
| Medication | -0.10 | 1.00 | -0.18 | 0.37 |
| Diagnosis (SZ vs BP) | 0.75 | 1.00 | 0.44 | 1.00 |
| Diagnosis (NAP vs AP) | 0.78 | 1.00 | -0.21 | 1.00 |

Either Pearson’s correlations (for continuous measures) or t tests (for categorical measures) were computed across all patients. No association was significant after FDR correction (*q*>0.05).

*Abbreviations: SZ=Schizophrenia spectrum disorders, including schizophrenia, schizoaffective disorder, schizophreniform, psychosis NOS, delusional disorder, or brief psychotic disorder. BP=Bipolar disorders with psychotic features, here for simplicity we also included major depressive disorder with psychotic features. NAP=Non-affective psychotic disorders, including schizophrenia, schizophreniform, psychosis NOS, delusional disorder, or brief psychotic disorder. AP=Affective psychotic disorders, including schizoaffective disorder, bipolar disorders with psychotic features, and major depressive disorder with psychotic features.*
